# The Out of East Asia model versus the African Eve model of modern human origins in light of ancient mtDNA findings

**DOI:** 10.1101/546234

**Authors:** Ye Zhang, Shi Huang

## Abstract

The first molecular model of modern human origins published in 1983 had the mtDNA phylogenetic tree rooted in Asia. This model was subsequently overlooked and superseded by the African Eve model in 1987 that was premised on the unrealistic infinite site assumption and the now failed molecular clock hypothesis. We have recently developed a new framework of molecular evolution, the maximum genetic diversity (MGD) hypothesis, which has in turn led us to discover a new model of modern human origins with the roots of uniparental DNAs placed in East Asia. While the African mtDNA Eve model has haplotype N as ancestral to R, our Asia model places R as the ancestor of all. We here examined ancient mtDNAs from the literature focusing on the relationship between N and R. The data showed that all three oldest mtDNAs were R with the 45000 year old Ust’-Ishim a basal type and the two ∼40000 year old samples sub-branch of R. Among the numerous mtDNAs of 39500-30000 year old, most were R subtype U and only two were N samples, the 39500 year old Oase1 and the 34425 year old Salkhit. These N types are basal and hence likely close to the root of N. These ancient DNA findings suggest that basal R is ∼5000 years older than basal N, thereby confirming the East Asia model and invalidating the African Eve model.

## Introduction

Investigation into the question of human origins has a long history and two competing models termed “Multiregional” and the “Recent Out-of-Africa” hypothesis have been proposed ^1^. In the Multiregional model ^2-4^, recent human evolution is seen as the product of the early and middle Pleistocene radiation of *Homo erectus* from Africa. Thereafter, local differentiation led to the establishment of regional populations which evolved to produce anatomically modern humans (AMH) in different regions of the world. *Homo* has been a single species since the genus first appeared in the fossil record ∼2.3-2.8 million years ago. The model has ample evidence from fossils and Paleolithic cultural remains but consistent molecular evidence has been lacking until very recently ^5^.

The single origin Recent Out of Africa model assumes that there was a relatively recent common ancestral population for *Homo sapiens* which already showed most if not all of the anatomical features shared by present day people. This population originated in Africa ∼200 ky ago, followed by an initiation of African regional differentiation, subsequent radiation from Africa, and final establishment of modern regional characteristics outside Africa ^1,6^. These modern Africans replaced the archaic *Homo* in Eurasia with limited genetic mixing ^7-11^. Support for this model comes from the African location of the earliest fossils of some AMH features ^12,13^ and the molecular clock and neutral theory interpretation of the greater genetic diversity in Africans ^6^.However, completely modern humans older than 20000 years have yet to be found in Africa but did exist in Daoxian South China, although the Daoxian site has no skeletons other than teeth, leaving it uncertain whether features other than teeth are also completely modern ^14,15^.

While long overlooked, there is in fact a third model based on mtDNA analyses. In 1983, researchers derived the first mtDNA phylogenetic tree and rooted the tree in Asia ^16^. Unfortunately, this Asia model was superseded 4 years later by the African Eve model without any valid reasons ^6^. The African Eve model assumes the molecular clock while the Asia model not. Given that the universal molecular clock is widely acknowledged to be unreal ^17-22^, a reexamination of these models is clearly warranted. Furthermore, the Africa model requires the neutral theory and the infinite site assumption, which, while largely sound as a null model and a framework for pre-saturation evolutionary processes, are not *a priori* valid and have met with great difficulties as an explanatory framework for most molecular evolutionary phenomena 23-27. Obviously, inferring human origins by using genetic diversity data must wait until one has a complete understanding of what genetic diversity means.

We have in recent years examined the longstanding genetic diversity riddle and developed a more complete evolutionary account, the maximum genetic diversity (MGD) hypothesis, that has been productive in addressing both evolutionary and biomedical problems ^21,28-36^. The MGD theory has solved the two major puzzles of genetic diversity, the genetic equidistance phenomenon and the much narrower range of genetic diversity relative to the large variation in population size ^27,28^. The genetic equidistance result of Margoliash in 1963 is in fact the first and best evidence for MGD rather than linear distance as mis-interpreted by the molecular clock and in turn the neutral theory ^19-21,28,34,37,38^. The MGD regards the genome to be mostly functional and under stabilizing selection. We have shown that genetic distances or diversities today are mostly at optimum saturation no longer related to time of evolution. However, the molecular clock and neutral theory misread reality of genetic diversity as still related to time of evolution.

Based on the MGD theory, we have developed the “slow clock” method that only uses slow evolving DNAs for demographic inferences because they are more likely to be neutral and away from the saturation phase of evolution ^5,34,39,40^. Our analyses have led to a new scheme of modern human origins largely consistent with the multiregional model in terms of autosomes where the first split among major human groups is placed at ∼1.9 million years ago ^5^. Different from the multiregional model, however, our new scheme has solved the origin problem of the modern uniparental DNAs that obviously can only have originated from a single region as the coalescence time of the modern mtDNA lineages is much shorter than 1.9 million years and as archaic humans, such as Neanderthals, Denisovans, and Heidelbergensis, clearly carried distinct lineages ^41^. Our model placed the roots of both mtDNA and Y chromosome in East Asia ^5^, which independently confirms the 1983 finding on the Asia origin of mtDNA ^16^. Our model furthermore specifically places the least differentiated haplotype R0 or R* as the ancestor of all mtDNA haplotypes, which is in contrast to the African Eve model that puts R downstream of haplotype N (Figure 1). The R0 haplotype is most common today in the Southern Chinese group in the 1000 genomes project, implicating the origin of the modern mtDNA lineage in Southern China ^5^. Our mtDNA tree was derived from analyzing present day haplotypes without needing to use ancient samples. Therefore, its validity can be independently tested by using ancient DNAs.

**Figure 1.**
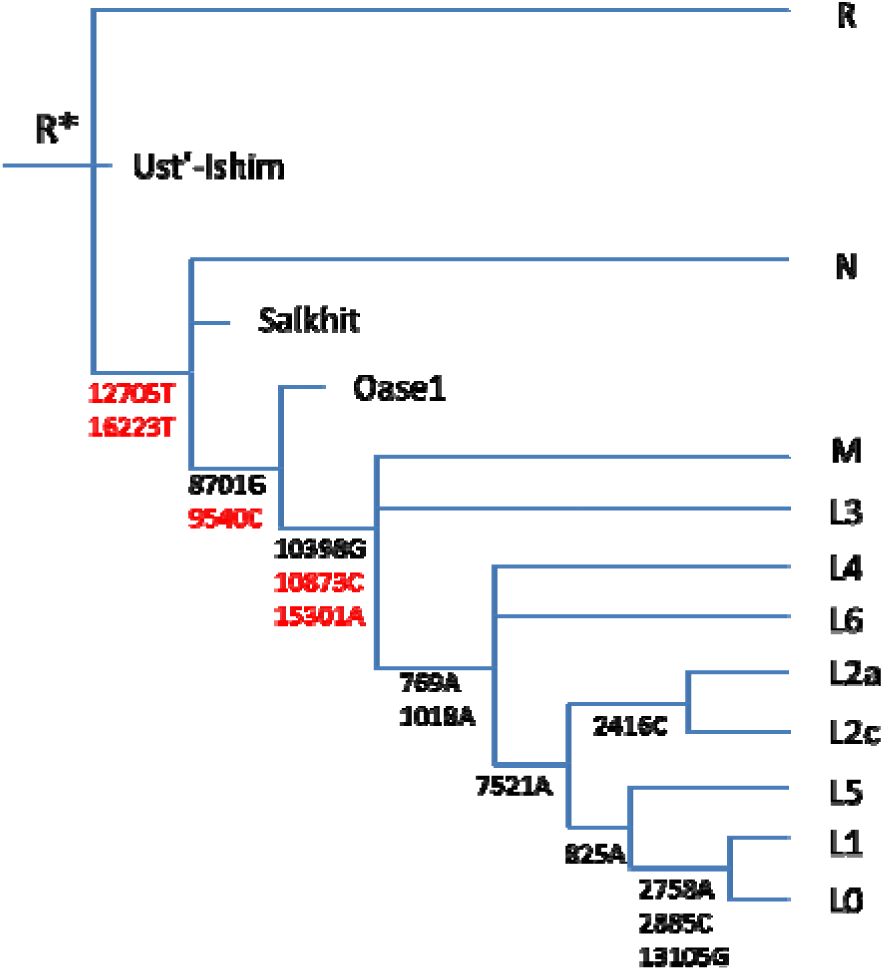
mtDNA phylogenetic tree of the Out of East Asia model. Only major braches and representative mutations are shown. Slow evolving sites (altering protein and RNA sequences) in black and fast sites in red. Also shown are several representative ancient DNAs that were placed based on their variant types. It should be noted that tree topology was built without making use of any ancient DNAs.

The Africa model assumes the infinite site assumption to classify haplotype relationships based on inference of ancestral and derived alleles with the allele identical to that of chimpanzees as the ancestral. However, the infinite site model is in fact unrealistic because there is overwhelming evidence of mutation saturation as explained by the MGD theory, which makes the inference of ancestral alleles meaningless. Actually, the Africa Eve phylogenetic tree cannot even meet the minimum standard of a scientific model, i.e., self-consistency, as it contains huge number of back mutations (reversions to an ancestral allele) that violate the infinite site assumption required to build the Africa model in the first place. These mutations can be found in the official mtDNA tree, phylotree (http://www.phylotree.org/), and are designated by an exclamation mark (!) following the position numbers in phylotree. The total number of these back mutations in phylotree is 1180 per our manual counting. For example, of the 5 defining mutations for haplotype N, one at site 15301 is a back mutation.

Another premise (inferred from the molecular clock) for the Africa model is that the original ancestor group should today show highest genetic diversity. However, as the MGD theory explains, reality is that different groups may show different maximum genetic diversity and human groups today are at saturation level of genetic diversity for the fast evolving DNAs ^5^. Africans do have the highest genetic diversity but only in fast changing variants and deleterious variants such as stopcodon gain/loss variants^5^. They in fact do have the highest genetic diversity in slow evolving neutral variants ^5^.

Unlike the Africa model, our Asia model, as well as that of Johnson et al ^16^, however, is based on the common sense that the original type should be the least differentiated. The original ancestor should mostly carry major alleles at SNP sites because as the ancestor population grew in size only a small fraction of the group should carry alternative mutant alleles (mutations are rare events). Indeed, the 45000 year old Ust’-Ishim’s mtDNA had fewer SNPs to begin with and the one he did have are all common alleles in today’s population (>67%) except one site 16150 ^10^. The Asia model also regards mutations in mtDNA to be functional, which has ample evidential support ^26,33,42,43^, and classifies haplotypes based on sharing of alleles with more weight on the slow evolving sites (altering protein or RNA sequences) ^5^. Hence, the Asia model is inherently more sound and self-consistent than the Africa model due to stronger theoretical foundations and far more realistic assumptions. Nonetheless, direct experimental test of these models is required to be absolutely certain. As our mtDNA tree was not derived by making use of ancient DNAs, it can be independently tested by using ancient samples. Recent discoveries in ancient DNAs have made such test possible and we here found the Asia model strongly supported by the ancient mtDNAs findings from the recent literature.

## Results and discussion

The relationship between N and R is the most obvious difference between the Africa and the Asia model. Of the three oldest mtDNAs, all turned out to be R, the 45000 year old Ust’-Ishim from Siberia ^10^, the 40328 year old Tianyuan from Northern China ^44^, and the 39805 year old Fumane2 from Riparo Bombrini Italy ^45^. Tianyuan was haplotype R subtype B as reported. Fumane2 was closest to extant HV or H haplotype (KP34013 and KF523402) per our analyses of the NCBI database. Only the oldest R sample Ust’-Ishim was found to be basal, consistent with expectations. All three had the expected mutations within R haplogroup (73G, 263G, 750G, 1438G, 2706G, 4769G, 7028T, 8860G, 11719A, 14766T and 15326G). But Ust’-Ishim had only 1 extra mutation (16150T), Tianyuan had 6 extra mutations (a 9 bp deletion, 5348, 5836, 11257, 16189, 16293), and Fumane2 had 3 (5585A, 16129A and 16304C). These extra mutations are mostly minor or rare alleles in present day people, indicating that Ust’-Ishim is more basal and less differentiated than the other two younger samples.

Among relatively younger samples of 39500-30000 years old, two N samples were found, the 39500 year old Oase1 from Romania ^9^ and the 34425 year old Salkhit from Mongolia ^46^. In contrast to the rarity of N, numerous samples of R subtype U were found in Europe as recently summarized by Deviese et al ^46^, indicating that R* basal samples may only be expected near the time when the root of R lived, which should be close to 45000 years ago as marked by the R* sample Ust’-Ishim. If R is indeed the root of modern mtDNA, then the modern lineage should not be much older than 45000 years old. In fact, while partially modern skeletons can be found in many sites in Africa and Asia at 310000-80000 years ago ^14,15,47-53^, no human skeletons that were fully modern for all key skeletal features have been found prior to 50000 years ago ^54^. As the demise of Neanderthals is thought to be associated with the arrival of modern humans and occurred at ∼40000 years ago ^45^, one can infer that the appearance of fully modern humans may not be much earlier than 40000 years ago. If it is much earlier, one would have the difficulty to explain the long lag before the demise of Neanderthals.

Both the N samples were found to be basal N* and not directly ancestral to any present day haplogroups ^46^. Oase1 had all N defining mutations except 8701G and 9540C and 6 extra mutations (3205A, 3462T, 4232C, 7158G, 8749C, 11016A). Salkhit had all N defining mutations and 10 extra mutations (4113A, 5492C, 8155A, 9456G slow, 9509C, 10364A, 14578T, 15077A slow, 16169T, 16092Gap). These extra mutations are also mostly minor or rare alleles in present day people. So, both of these N* samples appear to be far more differentiated than the R* type of Ust’-Ishim. As these two N* samples had ages of 39500-34425 years and no N subtypes were found in the numerous mtDNA samples from 40000-30000 years ago, which is in contrast to the abundantly found R subtype U, one can infer that the basal N* ancestor probably lived at 39500-34425 years ago and cannot be much earlier than 39500 years ago or older than R*. It is therefore highly unlikely to detect the presence of N in future samples older than 45000 years or the oldest R.

These two samples all have private mutations that are rarely found in today’s samples, indicating that those rare private mutations may represent adaptive changes related to adaptation to ancient environments and may not be used to classify these two samples as non-basal N. We note that Deviese et al ^46^ used the molecular clock method to calculate a split time between Salkhit and basal N* to be 19000 years and thus considered the appearance of N* basal type to be at ∼50000 years ago. However, such calculation may not be realistic as it did not consider the adaptive non-neutral roles of the ancient variants. Variations in mtDNA are mostly adaptive or under natural selection. Although distance among haplotypes may show some meaningful correlation with time of separation, such correlation are all derived by using present day haplotypes where the effects of variations due to adaption to environments from different eras are minimized. But comparing ancient DNA with present day haplotypes would represent a very different picture, because some variants in ancient DNAs may represent adaptive changes specific to the ancient environment. Those variants could be changed by today’s environments rather than by neutral drift. Therefore, one cannot use the molecular clock derived from present day samples to calculate the split time between an ancient sample such as Salkhit and other haplotypes of either today’s or ancient. In addition, if basal N* is indeed 50000 years old, it would contradict other findings. For example, one should find within the time window of 40000-30000 years ago differentiated N subtypes. And yet, only non-differentiated N were in fact found (Oase1 and Salkhit) that do not belong to any known present day haplotypes under N. In contrast, for a basal haplotype of truly 45000 years old such as R*,nearly all R samples found for the time window of 40000-30000 years ago were differentiated R subtypes such as U and B. Taken together, the combined data do not support placing basal N* at >40000 years ago.

There were two M samples aged 34000-35000 years, Goyet Q116-1 and Goyet Q376-3 from Belgium, found among samples older than 30000 years, and one M sample of 28000 years old from La Rochette France ^55,56^. As M is downstream of R and N in the Asia model but is parallel to N in the Africa model, the rarity of M relative to R and N in samples older than 35000 years is consistent with the Asia model but not the Africa model. Also, if the Africa tree is true where N and M are sister branches, the rarity of M in samples older than 35000 years ago further suggest that basal N* could not be 50000 years old.

That R* haplotype was found to be 5000 year older than N* haplotype as discussed above is consistent with the 2 SNP difference between R and N (Figure 1) and the mtDNA mutation rate of 2.67×10^-8^ substitution per site per year (5000 × 16500 × 2.67×10^-8^ = 2.2 substitution) as recently used by others ^46^. This suggests that the ages of the oldest R* and N* samples found so far were likely to be close to the real ages for the R and N ancestors. If the Africa model is true, the age of the original African L0 ancestor of all modern humans would be 120000 years old (35 SNPs separate L0 and N). However, as there are no completely modern humans found in Africa that are more than 20000 years old, the classification of L0 lineage as the original ancestor may be unrealistic. It is however placed as one of the last evolved in the Asia model, where its highly differentiated feature is a result of hybridization of AMH with archaic humans and coevolution of mtDNA with the admixed nuclear genome ^5^. It should be noted that the calculations here using molecular clock were all dealing with present day haplotypes within the same environmental or time period. Thus, effects of variants adaptive to environments of different eras are non-issues.

The original mtDNA type should leave more descendants today than the subtypes because mutations are rare events and occur only to rare individuals in a large population. We next asked whether the R haplotype is more popular today than other types by examining the data collected by the mitomap database (https://www.mitomap.org/MITOMAP). There are 64531 R samples in the database as inferred by using the R defining SNP 16223C, representing 53.6% of the whole collection. While the database may not be free of sampling bias and not all 16223C carrying samples are R due to independent mutations (again violating the infinite site assumption), one can nonetheless infer that it is highly likely for the R group to be larger than any other haplogroup as it is found larger than all the other haplogroups combined.

In contrast to the Asia model, the Africa model requires rare individuals with new mutations to have a reproductive advantage over ancestral populations, hence accounting for why the R group is much larger than any of its ancestor group, or why L0 is smaller than the mutant group L1’2’3’4’5’6, why L1 is smaller than the mutant group L2’3’4’5’6, why L5 is smaller than the mutant group L2’3’4’6, why L2 is smaller than the mutant group L3’4’6, why L6 is smaller than the mutant group L3’4, and why L4 is smaller than the mutant group L3. Such repeated reproductive advantages that must be granted to each round of new mutations leading to a new haplotype are in plain contradiction to the neutral mutation premise required to build the Africa model in the first place.

In summary, the Out of East Asia mtDNA model is inherently more sound and self-consistent than the African Eve model due to stronger theoretical foundations and far more realistic assumptions. The original R and N are likely to have ages close to those of the so far found oldest R* and N* samples, which showed that R* is older than N* by 5000 years. These results plainly invalidate the African Eve model and confirm the Out of Asia model. We expect our conclusion here to be further strengthened by future ancient DNA studies.

## Methods

Sequence alignment and mismatches were performed using blastn at the NCBI database. The number of back mutations in the mtDNA tree was inferred by counting the exclamation marks (!) that designate the back mutations in the official mtDNA tree phylotree (http://www.phylotree.org/). The number of R samples in the mitomap database https://www.mitomap.org/MITOMAP) were calculated by counting the number of haplotypes having the R defining SNP 16223C.

## Acknowledgments

Supported by the National Natural Science Foundation of China grant 81171880, the National Basic Research Program of China grant 2011CB51001 (S.H.).

## Additional Information

### Author contributions

Y.Z. and S.H. devised the project, analyzed the data and wrote the manuscript.

